# LincRNAs involved in DCS-induced fear extinction: Shedding light on the transcriptomic dark matter

**DOI:** 10.1101/834242

**Authors:** Stefanie Malan-Müller, Vladimir Barbosa C. de Souza, Willie MU Daniels, Soraya Seedat, Mark D. Robinson, Sîan M.J Hemmings

**Author notes:** Authors contributed equally to the manuscript (Stefanie Malan-Müller, Vladimir Barbosa C. de Souza).

## Abstract

There is a growing appreciation of the role of non-coding RNAs in the regulation of gene and protein expression. Long non-coding RNAs can modulate splicing by hybridizing with precursor messenger RNAs (pre-mRNAs) and influence RNA editing, mRNA stability, translation activation and microRNA-mRNA interactions by binding to mature mRNAs. LncRNAs are highly abundant in the brain and have been implicated in neurodevelopmental disorders. Long intergenic non-coding RNAs are the largest subclass of lncRNAs and play a crucial role in gene regulation. We used RNA sequencing and bioinformatic analyses to identify lincRNAs and their predicted mRNA targets associated with fear extinction that was induced by intra-hippocampally administered D-cycloserine in an animal model investigating the core phenotypes of PTSD. We identified 43 differentially expressed fear extinction related lincRNAs and 190 differentially expressed fear extinction related mRNAs. Eight of these lincRNAs were predicted to interact with and regulate 108 of these mRNAs and seven lincRNAs were predicted to interact with 22 of their pre-mRNA transcripts. On the basis of the functions of their target RNAs, we inferred that these lincRNAs bind to nucleotides, ribonucleotides and proteins and subsequently influence nervous system development, and morphology, immune system functioning, and are associated with nervous system and mental health disorders. Quantitative trait loci that overlapped with fear extinction related lincRNAs, included serum corticosterone level, neuroinflammation, anxiety, stress and despair related responses. This is the first study to identify lincRNAs and their RNA targets with a putative role in transcriptional regulation during fear extinction.

## Introduction

Posttraumatic stress disorder (PTSD) is a severe and debilitating disorder that is highly prevalent in individuals who experience one or more traumatic events (1). Dysfunctional fear extinction plays an integral role in the development of the disorder (2)(3). The development of PTSD involves a fear conditioning process, during which fear and anxiety responses are exaggerated and/or are resistant to extinction (4)(5)(6). During classical fear conditioning, a neutral (conditioned) stimulus (CS) is paired with an aversive (unconditioned) stimulus (US). Following adequate pairing of the CS and the US, the CS will eventually result in the same response as the US, and which is referred to as the conditioned response (CR). The CS subsequently has the ability to elicit a conditioned fear response, which can be triggered upon encountering a harmless stimulus associated with the trauma (7). In PTSD, the trauma is considered to be the US, and the conditioned fear response experienced by PTSD patients, even in the presence of seemingly harmless stimuli, is the CR (8)(9).

To relieve the anxiety and fear associated with the CS, deconditioning/desensitization to the learned fears, thus fear extinction, have to occur (10). Systematic desensitization to the CS relies on extinction and counterconditioning, two processes that involve learning. Exposure therapy is dependent on extinction learning to reduce the CR to stimuli that provoke anxiety and panic (11)(12). Recent research has indicated that fear extinction involves the formation of a new competing memory that inhibits the fear response, rather than deleting the original (traumatic) memory (13)(14). Treatment options for PTSD include exposure-based cognitive behavioural therapies (CBT) and pharmacological treatments, such as the selective serotonin reuptake inhibitors (SSRIs) (15) and serotonin-norepinephrine reuptake inhibitors (SNRIs) (16). Despite the relative efficacy of these treatments, a large number of PTSD patients do not respond optimally and/or relapse over time (17)(18)(19).

D-cycloserine (DCS), a partial N-methyl-D-aspartate (NMDA) receptor agonist at the glycine site on the NMDAR1 receptor subunit is effective in facilitating extinction learning in rats when administered before or immediately after extinction training (20)(21)(22). The coadministration of DCS and exposure-based CBT has also been proven to be effective in extinguishing fear in human trials of anxiety disorders (23)(24) and PTSD (25)(26). DCS administration facilitates generalized extinction of fear (21) and reduces the rate of relapse following successful exposure-based CBT (27). Studies have investigated the mechanisms of DCS facilitated fear extinction, with the majority focusing on either intra-amygdalar (28) or systemic administration (29)(30) and subsequent investigation of altered gene or protein expression (29). Memory consolidation, and by extension, fear extinction, requires dynamic gene and protein expression regulation; however, few studies have investigated transcriptional and post-transcriptional regulation during fear conditioning and fear extinction.

Only one-fifth of the human transcriptome is associated with protein-coding genes; noncoding RNAs (ncRNAs) are highly prevalent and outnumber coding genes (31). These ncRNAs therefore contribute significantly to the diversification of eukaryotic transcriptomes and proteomes. Currently, 172, 126 human lncRNA transcripts and 24, 879 rat lncRNA transcripts have been identified, encoded for by 96, 000 human and 22, 127 rat lncRNA genes (32). The majority of lncRNA genes are expressed in a cell-type-specific and developmental stage-specific manner (33). LncRNAs are categorized based on their proximity to proteincoding genes. The five categories are sense, antisense, bidirectional, intronic, and intergenic lncRNAs (34)(35). LncRNAs are involved in numerous sub-cellular processes, including cellular organelle formation and functions. Furthermore, lncRNAs are highly abundant in the central nervous system (CNS), and a vast number of neuronal lncRNAs are located adjacent to genes that encode transcriptional regulators and key drivers of neural development, including those involved in the regulation of neuronal differentiation (36), stem cell pluripotency (33), and synaptogenesis (37), implicating these lncRNAs in the regulation of these genes.

Involvement of lncRNAs in such a broad range of functions and processes is likely attributed to their ability to regulate transcription. LncRNAs can regulate the expression of neighbouring genes, both in-*cis* (38)(39) and -*trans* (40)(41) in several ways. One mechanism is promoter modifications, via histone modifications, nucleosome repositioning and DNA methylation, to either result in chromatin conformations accessible to transcription factors or by inhibiting the nuclear localisation of transcription factors (42), subsequently resulting in activation or repression of gene expression. LncRNAs also participate in RNA processing by hybridizing to mate RNA molecules, thereby influencing mRNA stability, RNA editing, pre-mRNA splicing, translation activation, or abolition of miRNA-induced repression (43). Furthermore, lncRNAs can interact at a protein level through physical interactions with alternative splicing regulators (44), and even act as scaffolds to arrange higher-order complexes, for instance during histone modification (42). Finally, lncRNAs have been implicated as signalling molecules during exosomal RNA transfer between cells, subsequently altering gene expression patterns in the recipient cell (45).

Long intergenic RNAs (lincRNAs) belong to a sub-class of lncRNAs that constitute more than half of lncRNA transcripts in humans (46). RNA sequencing of post-mortem brain samples from schizophrenia and bipolar disorder patients suggests the involvement of lincRNAs in mental disorders (47). A PTSD GWAS study conducted in African American women found a significant association with a novel RNA gene, lincRNA AC068718.1 (48). The authors hypothesised that this lincRNA, with predicted functions for telomere maintenance and immune function, may be a risk factor for PTSD in women. Their results add to emerging evidence that non-coding RNAs play a critical role in gene regulation and might be involved in the aetiology of stress-related disorders (49)(50). However, the modes of action and functions of most lncRNAs in disease remain to be elucidated.

LncRNAs have a rapid turnover rate, thereby providing lncRNAs with the ability to mediate rapid genomic responses to external stimuli, as opposed to the slow-acting response of protein-coding genes (51). LncRNAs of the CNS could, therefore, be involved in rapid cellular and molecular responses, such as those required for memory consolidation or extinction, making them attractive regulators to investigate in pathologies where memory processes are affected. The aim of this project was first, to identify lincRNAs associated with fear extinction as facilitated by the co-administration of behavioural fear extinction and intra-hippocampal DCS administration, in an animal model that simulated the core PTSD phenotypes, and second, to determine the role of these lincRNAs in regulating the transcriptome during fear extinction.

## Methods

### Animal model

All applicable international, national, and institutional guidelines for the care and use of animals were followed. All animal-related procedures were conducted in accordance with the ethical standards of Stellenbosch University’s Research Ethics Committee: Animal Care and Use (REC:ACU) (Ref: ACU/2010/006(A1)).

An adapted version of the PTSD animal model described by Siegmund and Wotjak (2007) was utilised (52). Briefly, 120 adult, male Sprague-Dawley rats were grouped into four experimental groups (30 rats per group) based on an associated fear conditioning paradigm using electric foot shocks. The groups received intrahippocampal administration of either DCS or saline: (1) fear-conditioned + intrahippocampal saline administration (FS), (2) fear-conditioned + intrahippocampal DCS administration (FD), (3) control + intrahippocampal saline administration (CS) and (4) control + intrahippocampal DCS administration (CD). Typical phenotypes associated with PTSD were assessed in this model (53), such as anxious/fearful behaviour (using the light/dark [L/D] avoidance test (54) and open field test (55)) and anhedonia (using the forced swim test (56)). The L/D avoidance test was found in our initial experiments to be the most sensitive behavioural test of anxiety and was subsequently used to differentiate maladapted (animals that displayed anxiety-like behaviour) from well-adapted (animals that did not display anxiety-like behaviour) sub-groups (refer to (57) for more methodological detail).

The following sub-groups are of interest to the current study: (i) control animals that received intra-hippocampal saline (CS, modelling a human control group); (ii) fear-conditioned animals that received intra-hippocampal saline (FS) and were maladapted (FSM, thus modelling a PTSD-like group), fear-conditioned animals that received intra-hippocampal DCS (FD) and were well-adapted (FDW, modelling a patient group exhibiting effective fear extinction due to treatment). We focussed on two sets of sub-group comparisons, namely the FSM vs. CS (modelling the fear conditioning process by comparing a PTSD-like group to controls) and FDW vs. FSM (modelling the fear extinction process by comparing a group that exhibit effective treatment-induced fear extinction to a PTSD-like group), and honed in on differentially expressed transcripts that were regulated in opposite directions in the two comparison groups. We, therefore, aimed to identify lincRNA and mRNA transcripts that were upregulated in response to fear conditioning in the FSM vs. CS group but downregulated during fear extinction in the FDW vs. FSM group, and *vice versa*, in order to identify lincRNAs and mRNAs specifically associated with the process of fear extinction induced by the co-administration of DCS and behavioural fear extinction (Fig. 1). These sets of opposite, differentially expressed lincRNAs and mRNAs will henceforth be referred to in this manuscript as the fear extinction related lincRNAs and fear extinction related mRNAs.

**Figure 1:**
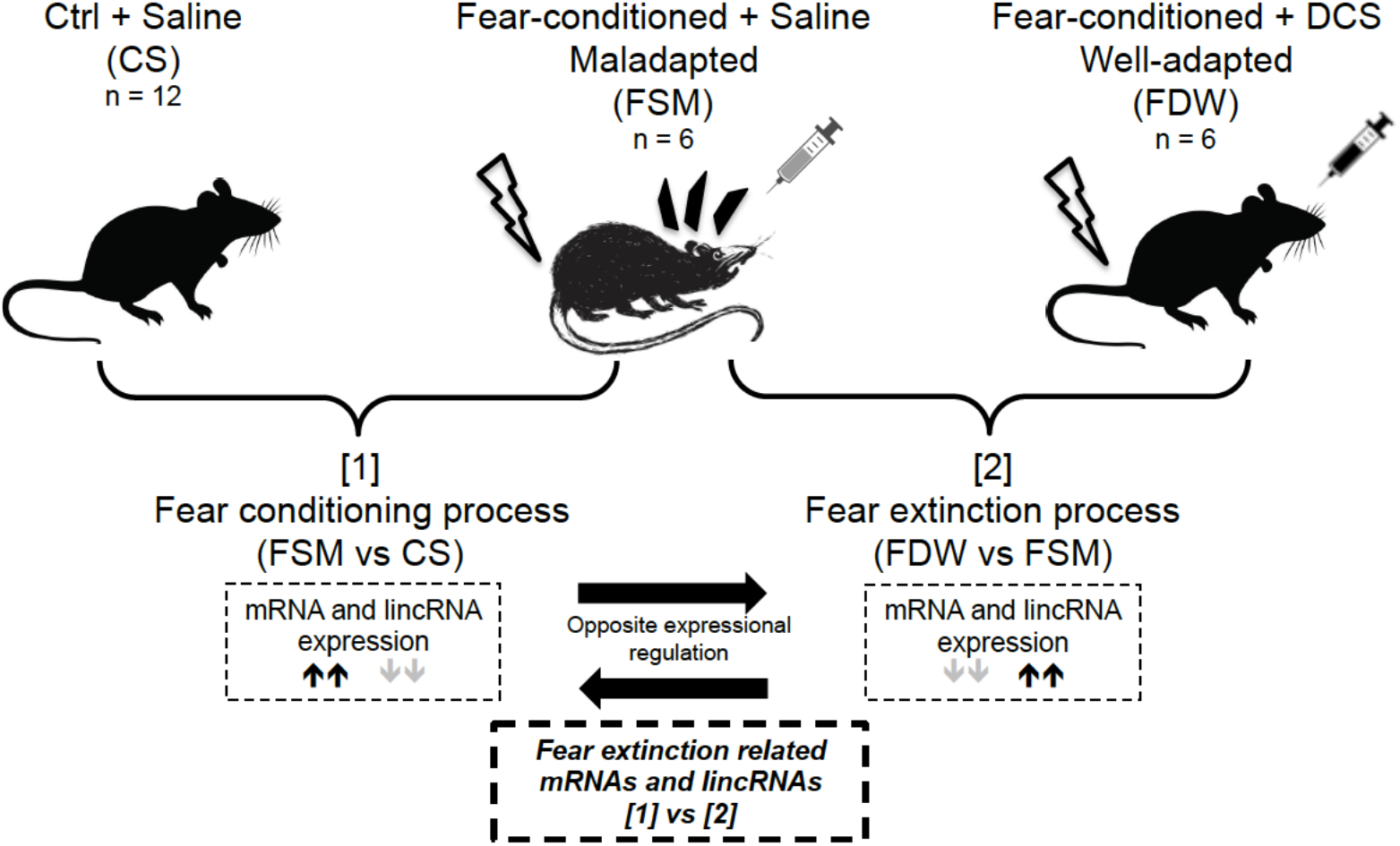
Diagram to explain animal sub-group comparisons utilised to identify fear extinction related mRNAs and lincRNAs following the co-administration of DCS and behavioural fear extinction

### RNA extraction and sequencing

RNA was extracted from the left dorsal hippocampal (LDH) regions of 30 rats (six rats each per FSM, FDW, FSM sub-groups and 12 rats in the CS sub-group) using the RNeasy Plus Mini Kit (Qiagen, Hilden Germany). RNA extraction, quantification and sequencing were performed on the 30 LDH RNA samples as described in (57).

### Bioinformatics and statistical analyses

FASTQC was used for quality assessment of RNA sequencing data. To identify differentially expressed transcripts for the sub-group comparisons FSM vs. CS and FDW vs. FSM, expression was quantified using *Salmon* (version 0.8.2) (58) with the Ensembl release-87 catalogue (coding and non-coding transcripts) (59) and *tximport* (version 1.12.0) (60) was used to import transcript counts into R v3.5.1 (61). We used the *edgeR* package (62) to identify differentially expressed lincRNAs and coding RNAs. The robust generalized linear model approach, as described by Zhou et al., (2014)(63), was used to estimate the dispersion parameter and make inferences for changes in expression.

### *In silico* prediction of lincRNA-mediated gene expression regulation during fear extinction

LincRNAs perform their diverse functions by interacting with a range of molecules, of which RNAs appears to be favoured (64). Therefore, the identification of potential mRNA targets of lincRNAs can help us better understand the functions of lincRNAs and determine how they regulate the transcriptome to facilitate fear extinction. The LncTar tool, developed for large-scale predictions of RNA-RNA interactions (65), was used to identify potential lincRNA– mRNA and lincRNA-pre-mRNA interactions within the sets of fear extinction related lincRNAs and mRNAs. LncTar uses base pairing and determines the minimum free energy joint structure of the two RNA molecules (65). The fasta sequences of fear extinction related lincRNA, mRNA and pre-mRNA transcripts were sourced from ensemble.org and used as the input files. The normalized free energy (ndG) cutoff, which indicates the relative stability of internal base pairs in the paired RNAs (66)(67)(68), was set to the second-highest stringency of - 0.15, to limit the results to interactions with a high probability.

### Identifying possible functions of fear extinction related lincRNAs

Although many lincRNAs have been identified, little is known about their functions. The functions of lincRNAs can be deduced from their genomic location or the functions of their targets. To facilitate biological interpretation of large sets of differentially expressed transcripts, gene set enrichment analyses were used to group transcripts together based on their functional similarity (69). To glean information about the functions of the fear extinction related lincRNAs, we investigated the biological processes, molecular functions and pathways (using Comparative Toxicogenomics Database (CTD) (http://ctdbase.org/tools/analyzer.go)) (70) and diseases (Rat Genome Database (RGD) (https://rgd.mcw.edu/rgdweb/enrichment/start.html) (71) associated with the predicted interacting fear extinction related mRNAs. Biological processes and molecular function categories were considered overrepresented if the Bonferroni-corrected p-value was < 0.01, and for pathways when Bonferroni-corrected p-value was < 0.05. Only the higher-order parental or ancestral terms for enriched diseases and higher GO levels for biological processes (levels 4 - 6) and molecular functions (levels 3 - 5) will be reported to simplify results and highlight key findings.

### LincRNA quantitative trait loci (QTL) overlap

The genomic locations of the fear extinction related lincRNAs were also inspected to determine proximity to QTLs (genes or genomic loci that contribute significantly to the variation in phenotypes/traits (72)), which could suggest putative functions of these lincRNAs (73)(74)(75). RGD (71) was used to identify corresponding RGD names of the fear extinction related lincRNAs and was subsequently used to identify QTLs that overlap with the location of these lincRNAs.

## Results

### RNA sequencing and differential expression analysis

RNAs with FDR < 0.05 and absolute log-fold-change (logFC) ≥ 1 are illustrated as red points on the minus-add (MA) plot of log-fold-change versus log-counts-per-millions (Fig. 2A). The overlap of detected differentially expressed features were calculated and plotted using the *UpSet* (76) package. An UpSet plot is a visualization approach for the quantitative analysis of sets, their intersections and aggregates of intersections, and serves as an alternative to Venn diagrams (Fig. 2B).

**Figure 2:**
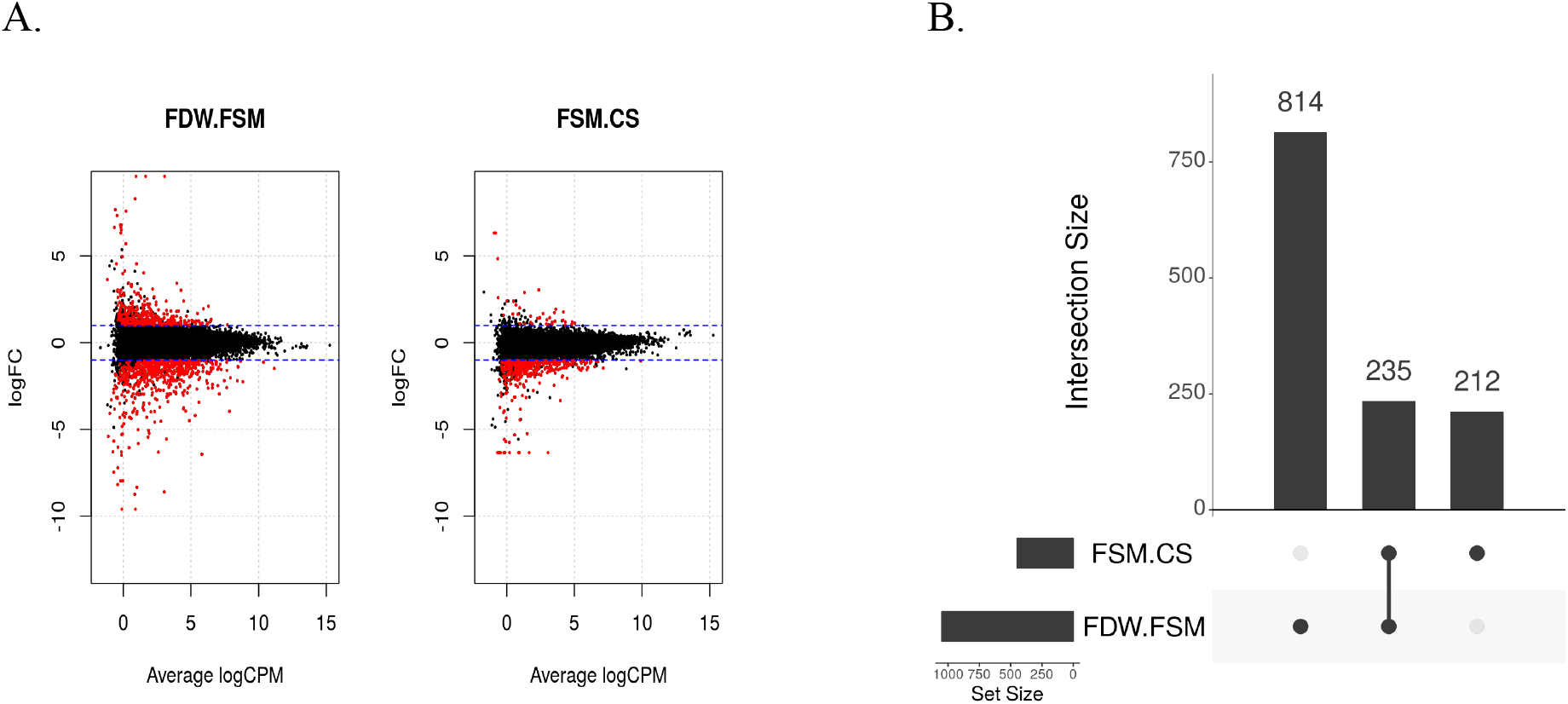
**(A)** MA (minus-add) plots of log-fold-change (logFC) versus average log-counts-per-millions (logCPM)/ average abundance of lincRNA and mRNA transcripts. The blue dotted lines indicate logFC cut off values of > 1 or <-1. Red points are significantly differentially expressed transcripts (including lincRNAs and mRNAs) at a false discovery rate (FDR) of 5%. **(B)** An UpSet plot illustrating unique and shared differentially expressed lincRNA and mRNA transcripts between the two sub-group comparisons, FSM vs. CS and FDW vs. FSM.

A total of three transcripts were up-regulated in the FSM vs. CS and down-regulated in FDW vs. FSM and a total of 230 transcripts were down-regulated in FSM vs. CS and up-regulated in FDW vs. FSM (Supplementary Tables 1-5). One transcript was down-regulated in both these subgroup comparisons and another transcript was up-regulated in both subgroup comparisons, resulting in a final sum of 235 overlapping transcripts (Fig. 2B) (Supplementary Figures 1-2 show all differentially expressed mRNA and lincRNA transcripts for the two comparison groups). A total of 190 fear extinction related mRNA transcripts were regulated in opposite directions between the two subgroup comparisons of interest ([1] FSM vs. CS and [2] FDW vs. FSM), with three transcripts up-regulated and 187 transcripts down-regulated in the fear conditioning comparison group [1] relative to the fear extinction comparison group [2] (Fig. 3a) (Supplementary Table 6) and 43 lincRNA transcripts were down-regulated in the fear conditioning comparison group [1] relative to the fear extinction comparison group [2] (Fig. 3b) (Supplementary Table 7). A breakdown of the differentially expressed lincRNA and mRNA transcripts is provided in Table 1 (Supplementary Figures 1-2, Supplementary Tables 2-7).

**Figure 3:**
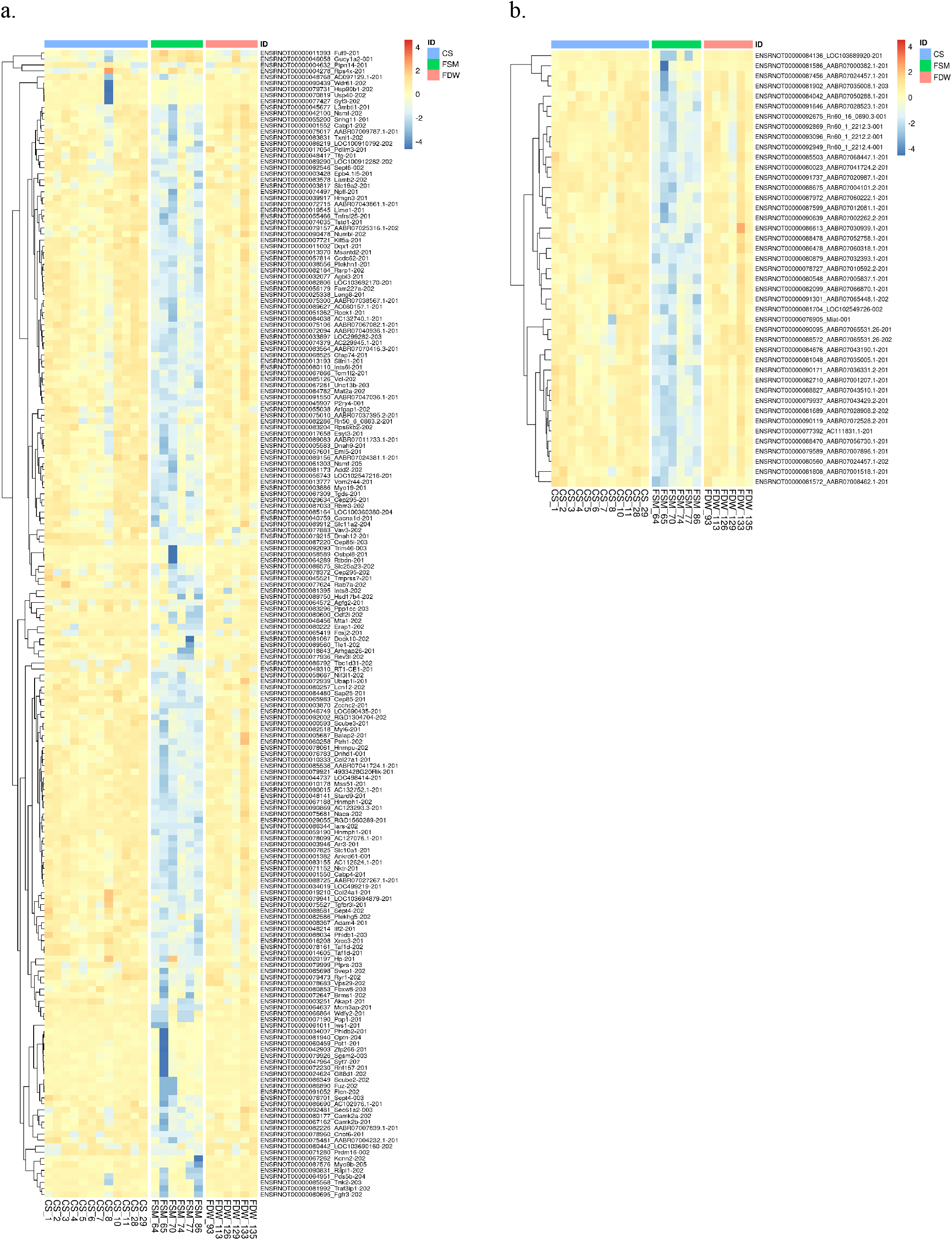
Heatmaps of a) fear extinction related mRNA transcripts and b) fear extinction related lincRNA transcripts that were differentially expressed between [1] FSM vs. CS and [2] FDW vs. FSM

**Table 1:**
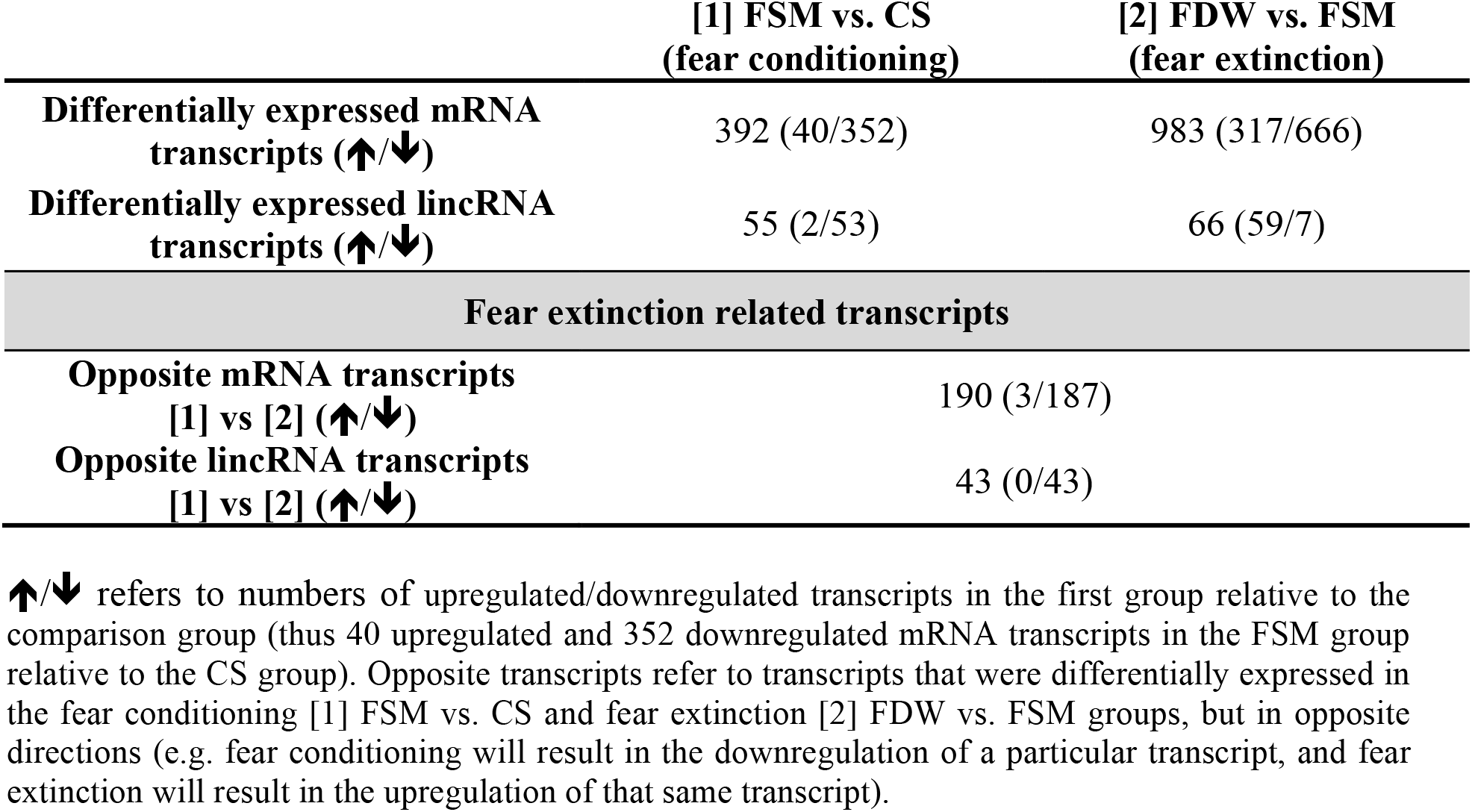
Summary of the number of differentially expressed mRNA and lincRNA transcripts for CS vs. FSM and FSM vs. FDW sub-groups

|  | [1] FSM vs. CS<br>(fear conditioning) | [2] FDW vs. FSM<br>(fear extinction) |
| --- | --- | --- |
| **Differentially expressed mRNA transcripts (↑/↓)** | 392 (40/352) | 983 (317/666) |
| **Differentially expressed lincRNA transcripts (↑/↓)** | 55 (2/53) | 66 (59/7) |
| **Fear extinction related transcripts** |  |  |
| **Opposite mRNA transcripts**<br>**[1] vs [2] (↑/↓)** | 190 (3/187) |  |
| **Opposite lincRNA transcripts**<br>**[1] vs [2] (↑/↓)** | 43 (0/43) |  |

↑/↓ refers to numbers of upregulated/downregulated transcripts in the first group relative to the comparison group (thus 40 upregulated and 352 downregulated mRNA transcripts in the FSM group relative to the CS group). Opposite transcripts refer to transcripts that were differentially expressed in the fear conditioning [1] FSM vs. CS and fear extinction [2] FDW vs. FSM groups, but in opposite directions (e.g. fear conditioning will result in the downregulation of a particular transcript, and fear extinction will result in the upregulation of that same transcript).

### *In silico* prediction of lincRNA-mediated gene expression regulation during fear extinction

LncTar predicted 119 lincRNA-mRNA interactions (Supplementary Table 8), from the 43 fear extinction related lincRNA transcripts, eight lincRNAs were predicted to interact with 108 fear extinction related mRNAs (Fig. 4). There were nine mRNA transcripts that were targeted by more than one lincRNA and the lincRNA ENSRNOT00000076905 had the highest number of predicted mRNA interactions, yielding 89 in total.

**Figure 4:**
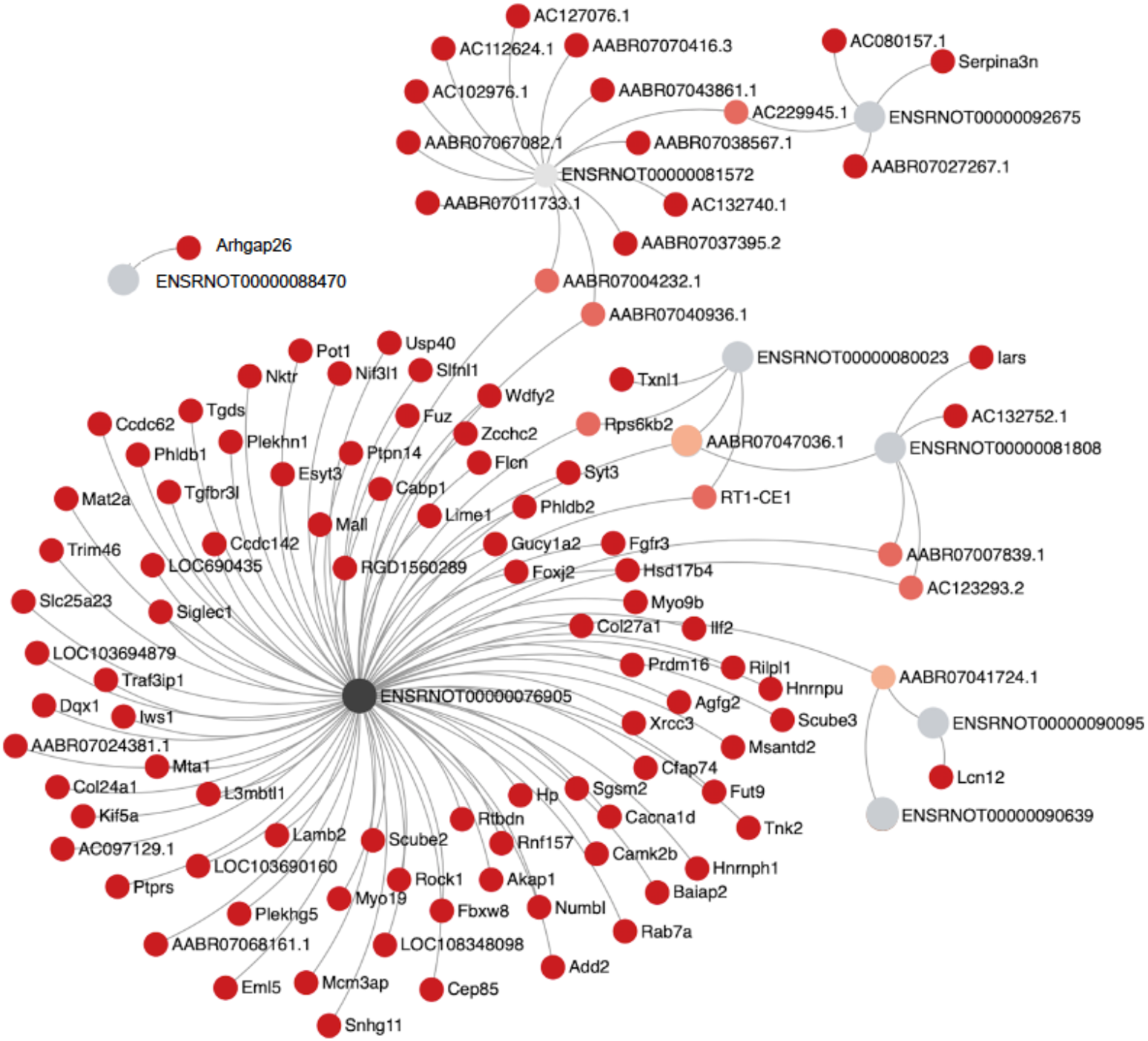
Fear extinction related lincRNA-mRNA transcript interactions. LncTar predicted lincRNA-mRNA transcript interactions; grey circles represent the eight fear extinction related lincRNA transcripts predicted to interact with the fear extinction related 119 mRNA transcripts (red and orange circles). Orange circles represent the nine mRNA transcripts that interacted with more than one lincRNA transcript.

LncTar predicted 30 interactions between differentially expressed lincRNAs and pre-mRNAs, with seven lincRNAs predicted to interact with 22 pre-mRNA transcripts (Fig. 5, Supplementary Table 9). There were six pre-mRNA transcripts for which there was no corresponding interaction between the lincRNA and its mature mRNA transcript (Table 2, indicated with stars in Figure 5). This is likely a result of the interaction regions falling within intronic regions or over exon-intron boundaries, therefore representing lincRNA-pre-mRNA interactions with possible splicing effects.

**Figure 5:**
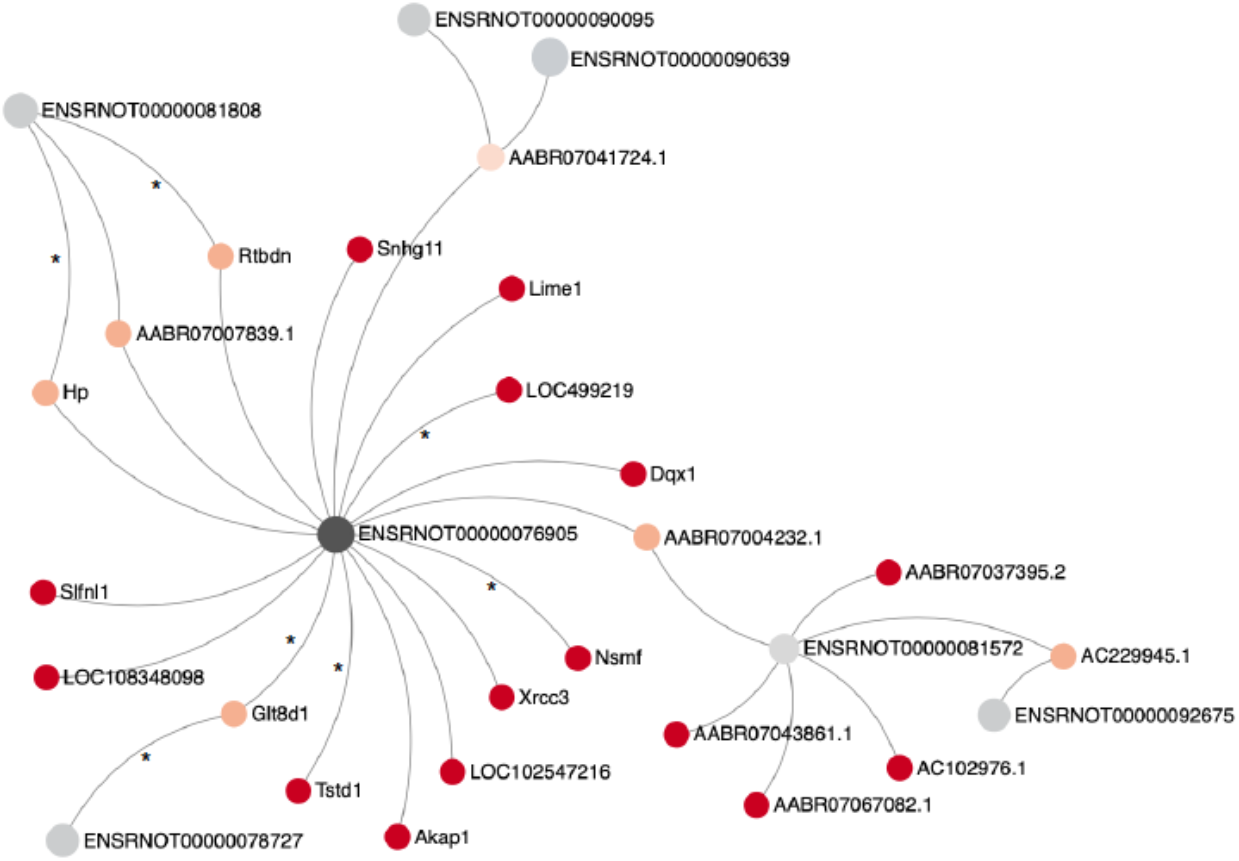
Fear extinction related lincRNA-pre-mRNA transcript interactions. LncTar predicted lincRNA-mRNA transcript interactions; grey circles represent the seven fear extinction related lincRNA transcripts predicted to interact with the 22 fear extinction related pre-mRNA transcripts (red and orange circles). Orange circles represent the nine mRNA transcripts that interacted with more than one lincRNA transcript. Stars indicate the six pre-mRNA transcripts and seven interactions for which there was no corresponding interaction between the lincRNA and its mature mRNA transcript

**Table 2:** Predicted fear extinction related lincRNAs and pre-mRNAs interactions that may affect splicing

| pre-mRNA target | LincRNA | Target region |
| --- | --- | --- |
| *Glt8d1* | ENSRNOT00000076905 | intron2 |
| *Glt8d1* | ENSRNOT00000078727 | intron 2, exon3, intron 3 |
| *Hp* | ENSRNOT00000081808 | intron 2, exon 3, intron 3 |
| *Nsmf* | ENSRNOT00000076905 | intron4 |
| *LOC499219* | ENSRNOT00000076905 | intron1 |
| *Rtbdn* | ENSRNOT00000081808 | intron1 |
| *Tstd1* | ENSRNOT00000076905 | intron 3 to exon 4 |

*Glt8d1* - glycosyltransferase 8 domain containing 1 gene, *Hp* – haptoglobin gene, *Nsmf* - NMDA receptor synaptonuclear signaling and neuronal migration factor gene, *LOC499219* – gene transcribing hypothetical protein, *Rtbdn* - retbindin gene, *Tstd1* - thiosulfate sulfurtransferase like domain containing 1 gene, UTR – untranslated region

### Gene ontology, disease and pathway enrichment analyses to predict functions of fear extinction related lincRNAs

A total of 81 lincRNA-interacting fear extinction related mRNAs were used in the enrichment analyses, 27 transcripts were clone-based transcripts with unknown functions and were excluded by CTD and RGD databases. Figure 6 shows the most enriched disease (top 25) (Fig. 6a), biological process (Fig. 6b) and molecular function terms (Fig. 6c), based on the Bonferroni corrected p-values and the number of annotated genes for each term (Supplementary Tables 10 – 12 contain exact p-values and all mRNA transcripts associated with each term).

**Figure 6:**
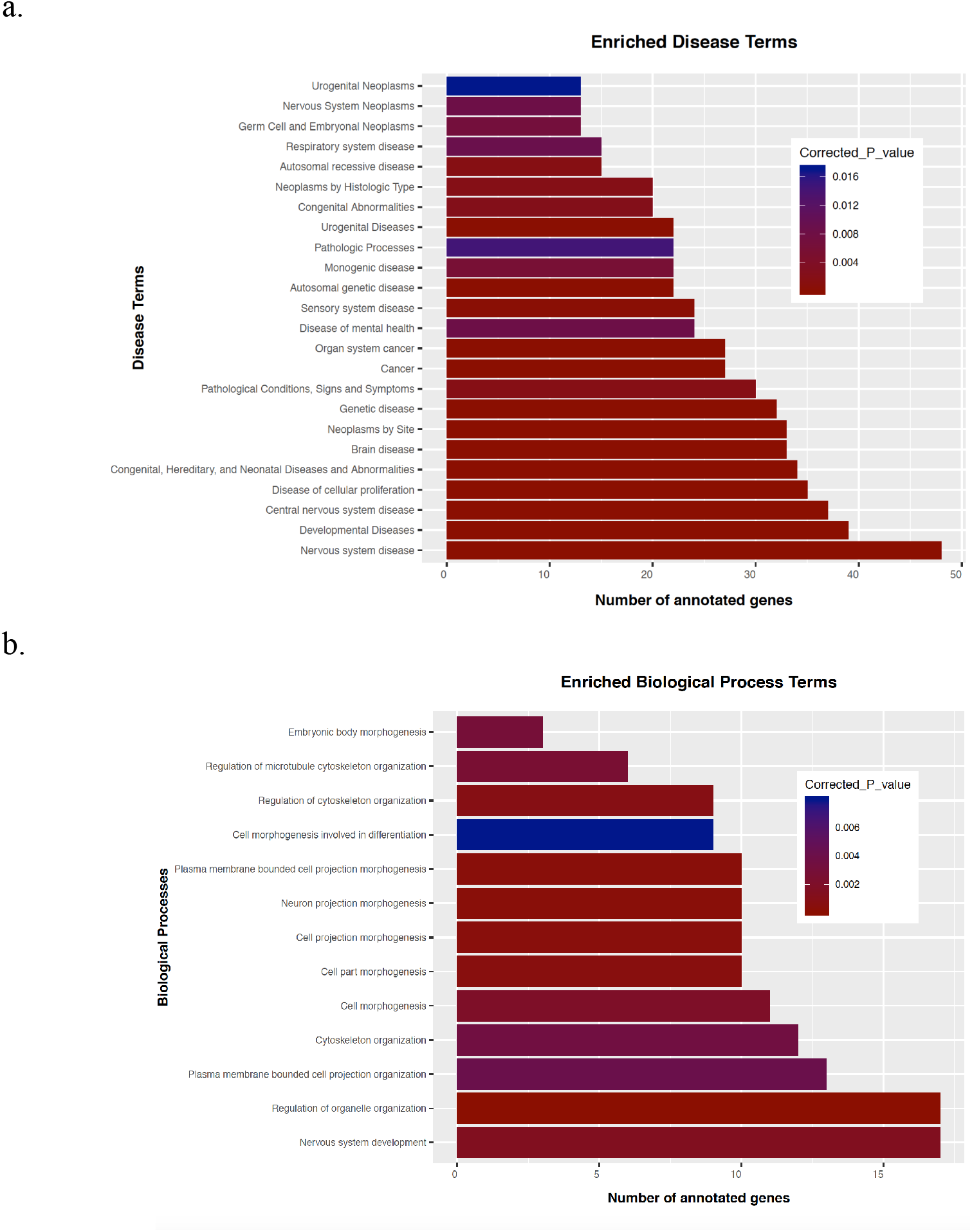
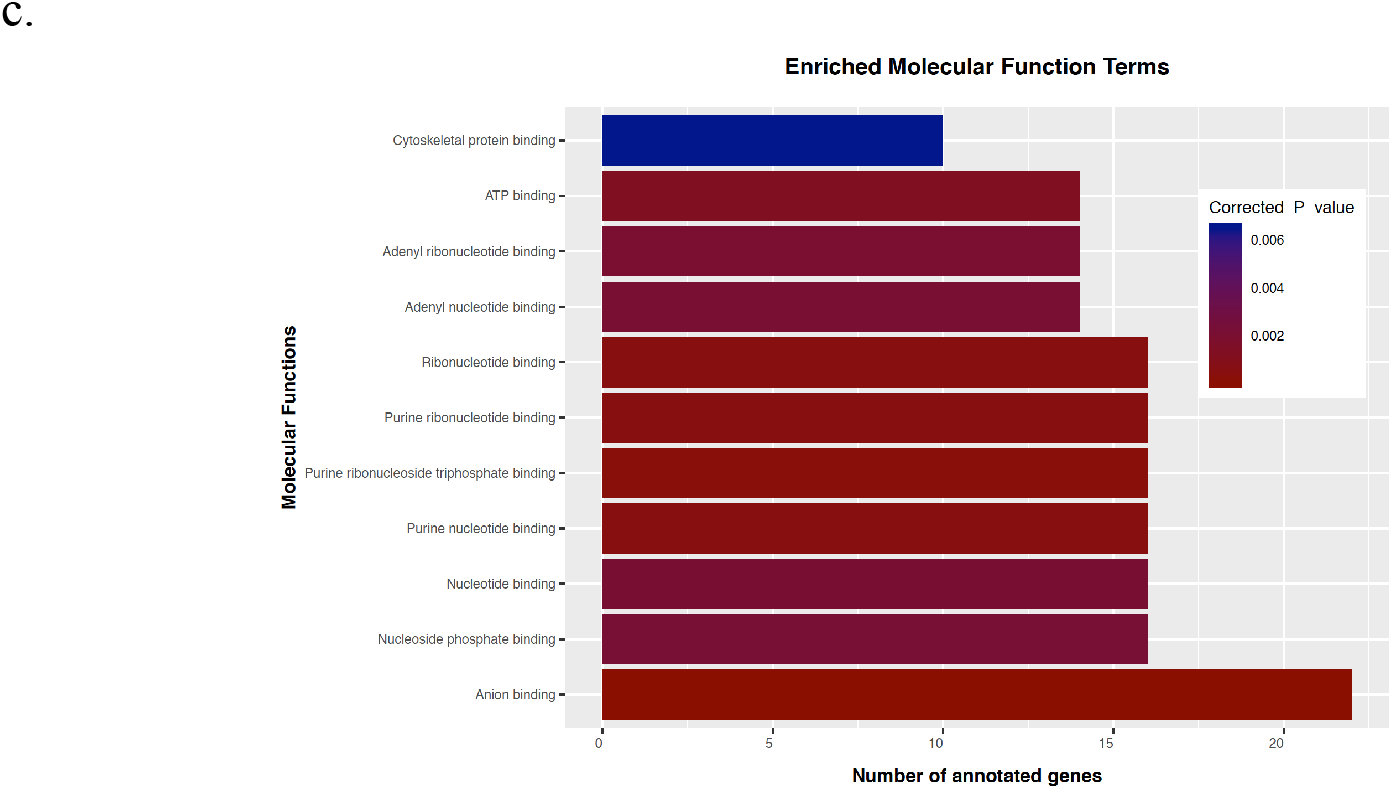
(a) Enriched disease terms (top 25), (b) biological processes and (c) molecular functions associated with the 61 fear extinction related mRNAs predicted to interact with fear extinction related lincRNAs. Bars are filled according to the significance level of Bonferroni corrected p-values; significance increases from blue to red.

A variety of disease terms were associated with these fear extinction related lincRNAs (Fig. 6a); *Nervous system disease* was the most significant term and several related terms were also enriched for, such as *Brain disease*, *Central nervous system disease* and *Disease of mental health*. A total of 48 mRNA transcripts were associated with these disease terms, of which the lincRNA ENSRNOT00000076905 was predicted to interact with 45 mRNA transcripts. Additional predicted interactions included ENSRNOT00000088470 with *Arhgap26*, ENSRNOT00000080023 with *Rps6kb2* and ENSRNOT00000092675 with *Serpina3n*.

Of the 13 enriched biological process terms, *Neuron projection morphogenesis* and *Nervous system development* were of particular interest (Fig. 6b). A total of 17 mRNA transcripts were associated with these terms, of which the lincRNA ENSRNOT00000076905 interacted with 15 neurogenesis-associated mRNA transcripts (*Add2, Baiap2, Camk2B, Fbxw8, Fuz, Lamb2, Nif3L1, Numbl, Prdm16, Ptprs, Rnf157, Rock1, Syt3, Traf3Ip1* and *Trim46*), ENSRNOT00000088470 interacted with *Arhgap26* and ENSRNOT00000076905 interacted with *Baiap2*. Eleven molecular function terms were associated with the lincRNA-interacting mRNA transcripts (Fig. 6c), with the main molecular functions encompassed in the broader terms of nucleotide, ribonucleotide and protein binding. A total of 26 mRNA transcripts were associated with these terms and 24 of these transcripts were predicted to interact with ENSRNOT00000076905. Furthermore, ENSRNOT00000080023 interacted with *Rps6kb2* and ENSRNOT00000088470 interacted with *Arhgap26* (Fig. 5). Two pathways, namely *Immune system* and *Signalling by Rho GTPases*, were associated with 15 of the fear extinction related mRNAs that interacted with the fear extinction related lincRNAs (Supplementary Table 13). Fourteen of these transcripts were predicted to interact with ENSRNOT00000076905; ENSRNOT00000080023 interacted with *Rps6kb2* and ENSRNOT00000088470 interacted with *Arhgap26* (Fig. 5). The *Oxytocin signalling pathway* was associated with four of the fear extinction related mRNAs, however, the association was not statistically significant (adjusted p = 0.068).

### LincRNA quantitative trait loci (QTL) overlap

To infer possible roles of the fear extinction related lincRNAs that may have regulated the transcription of genes in close proximity; we identified QTLs that overlap with their genomic locations. Four of the 43 fear extinction related lincRNAs had available corresponding RGD names to use in the RGD QTL overlap analysis. Table 3 summarises the most significant (LOD ≥ 3, p-value < 0.01) and relevant (in the context of fear extinction) QTLs.

**Table 3:**
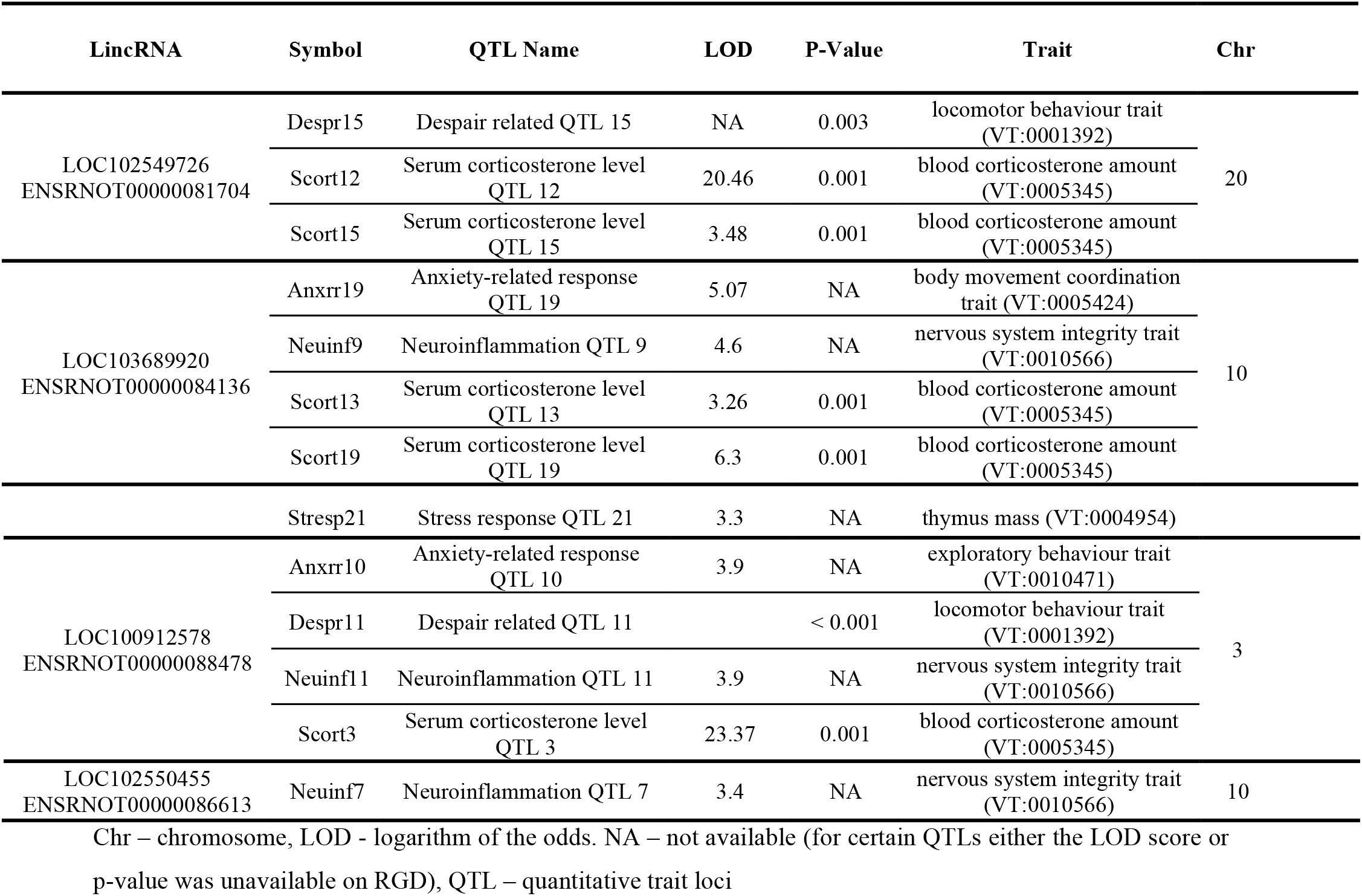
LincRNAs that overlap with QTLs of interest

## Discussion

This study aimed to identify lincRNAs that might be involved in the molecular mechanisms of DCS-facilitated fear extinction. LincRNAs associated with fear conditioning were identified as differentially expressed lincRNAs in FSM vs. CS, and those associated with fear extinction were differentially expressed in FDW vs. FSM, and in the opposite direction as in the FSM vs. CS group, and were referred to as fear extinction related lincRNAs. To determine the functions of these lincRNAs, we identified differentially expressed fear extinction related mRNAs, and used *in silico* prediction software to determine if these lincRNAs may have regulated the expression of these mRNAs or their precursor pre-mRNAs through RNA-RNA hybridisation complexes. Gene ontology enrichment analyses were performed for these targetted mRNAs to identify associated diseases, biological processes, molecular functions and pathways associated with the fear extinction related lincRNAs that targetted these mRNAs. In addition, we identified lincRNAs whose genomic location overlapped with QTLs that could explain why these lincRNAs may be involved in the process of DCS-facilitated fear extinction.

Our prediction analyses indicated that, based on sequence homology, eight lincRNAs could interact with 108 mature mRNA transcripts. These interactions may have influenced RNA editing, mRNA stability, translation activation and miRNA-mRNA interactions of genes that are important for fear extinction (see gene ontology discussion). Seven lincRNAs were predicted to interact with 22 pre-mRNA transcripts. Six of these interactions were not predicted for the corresponding mature mRNA transcripts, where the hybridization occurred in intronic regions or within exon-intron boundaries. We hypothesise that these interactions may influence translation and splicing events in those transcripts. Therefore, the differential expression of some of the fear extinction related mRNAs could be attributed to alternative splicing of their fear extinction related pre-mRNAs.

Of particular interest was the interaction between ENSRNOT00000076905 and the pre-mRNA of the NMDA receptor synaptonuclear signalling and neuronal migration factor gene (*Nsmf*), since DCS is a partial NMDAR agonist and binding of DCS to NMDARs facilitates extinction learning (20)(21)(22). One study found that an *Nsmf* knockout murine model, deficient for the Jacob protein transcribed by *Nsmf*, exhibited hippocampal dysplasia, impaired BDNF-signaling during dendritogenesis, and phenotypes related to the lack of BDNF-induced nuclear import of Jacob (which is NMDAR-dependent). The authors proposed a role for the Jacob protein in hippocampal dendrite-and synaptogenesis (77). Our data indicated that *Nsmf* was downregulated during fear conditioning (FSM vs. CS), and upregulated during DCS-facilitated fear extinction (FDW vs. FSM), potentially through the activation of NMDARs by DCS, which facilitates nuclear import of the Jacob protein. This could promote dendrite-and synaptogenesis, and possibly facilitate fear extinction. Furthermore, the *Nsmf* gene undergoes extensive splicing, with more than 20 known splice isoforms. The overexpression of one such splice isoform (*Δex9-Jacob*) in primary neurons, resulted in decreased dendritic complexity and number of synapses (78), emphasizing the importance of Jacob splice variants in hippocampal synaptogenesis, a process central to learning and memory (79)(80). Our analysis predicted an interaction between ENSRNOT00000076905 and *Nsmf* pre-mRNA, which may have resulted in alternative splicing, and alternative isoforms of the Jacob protein, with possible implications for hippocampal synaptogenesis and, possibly, fear extinction.

To predict the functions of fear extinction related lincRNAs, we performed gene ontology, and disease and pathway enrichment analyses on the set of predicted fear extinction related mRNA targets (therefore all interactions referred to in the enrichment analyses are fear extinction related transcripts, therefore differentially expressed and in the opposite direction between [1] FSM vs. CS [2] FDW vs. FSM). The most enriched disease term was *Central Nervous system disease*, but other synonymous disease terms were also significant, including *Disease of mental health*. The lincRNA ENSRNOT00000076905 was predicted to interact with the majority of mRNAs enriched in these disease terms (Supplementary Table 10). The likely reason for the vast number of predicted interactions of this lincRNA is its short length (140bp), which increases the likelihood of complementary hybridization to several mRNA transcripts. Additional lincRNAs predicted to interact with mRNAs enriched for central nervous system disease terms, included ENSRNOT00000088470, ENSRNOT00000080023 and ENSRNOT00000092675. These lincRNAs could, therefore, be involved in diseases that affect the CNS and mental health, by targeting and regulating genes associated with these disease terms.

For biological process enrichment, one lincRNA, ENSRNOT00000076905, was predicted to interact with 15 of the 17 mRNA transcripts involved in nervous system development and neuronal projection (neurogenesis). We hypothesise that this lincRNA is involved in neurogenesis, neuronal projection and extension. The fear extinction protocol consisted of reexposure to the shock chamber (without shock application), together with intra-hippocampal DCS administration. Our findings, therefore, suggest that DCS facilitated the process of fear extinction by promoting hippocampal neurogenesis. This correlates with earlier findings reporting that hippocampal DCS infusion increased neuronal proliferation and neural plasticity mediated by hippocampal NMDA receptors, which promoted the acquisition and retrieval of extinction memory (81). These results shed further light on the molecular mechanisms behind DCS-facilitated fear extinction, where the lincRNA ENSRNOT00000076905 may interact and regulate the expression of several mRNA transcripts, to ultimately facilitate fear extinction via neurogenesis.

Other predicted lincRNA interactions with mRNAs enriched for the biological process neurogenesis, include that of the brain-specific angiogenesis inhibitor 1-associated protein 2 (*Baiap2*) with ENSRNOT00000076905 and Rho GTPase activating protein 26 (GTPase Regulator) (*Arhgap26*) with ENSRNOT00000088470. The *Baiap2* gene encodes a synaptic protein whose hippocampal expression is required for learning, memory (82) and social competence (83). Furthermore, a SNP in *BAIAP2* has been associated with negative modulation of memory strength in humans (84), a process that plays an important role in PTSD (85). A study that investigated early-life programming and related gene x environment interactions in the context of anxiety and depression, found that *Baiap2* was downregulated following prenatal stress exposure (86). We, therefore, hypothesise that DCS administration reversed the negative effect that fear conditioning had on the expression of *Baiap2*, via a proposed ENSRNOT00000076905-mediated upregulation of *Baiap2*, thereby promoting fear extinction learning (87).

The *Arhgap26* gene transcribes a protein that is part of the Rho family of GTPases and interestingly, *Signalling by Rho GTPases* was one of the enriched pathways associated with fear extinction related lincRNAs. This pathway has been implicated in the regulation of learning and memory (88). A proposed mechanism underlying memory formation is the rearrangement of synaptic connections in neural networks. Dendritic spines receive the majority of excitatory synapses (89)(90) and undergo dynamic, experience-dependent changes (91). Furthermore, changes in dendritic spine morphology have been observed during long-term potentiation (LTP), a process that models the activity-dependent changes of synaptic efficacy and the cellular basis of learning (92)(93). Dendritic spine morphology and rearrangement are controlled by the neuronal actin cytoskeleton (94)(95), of which actin assembly, polymerization and actomyosin contraction are mainly regulated by small GTPases of the Rho family (96)(97)(98). LTP induction is associated with actin cytoskeletal reorganization, which is characterized by a sustained increase in F-actin content within dendritic spines. This increased F-actin content is dependent on NMDA receptor activation and involves the inactivation of actin-depolymerizing factor (cofilin) (94). It is thus possible that DCS activated the NMDA receptor, resulting in increased F-actin content and subsequent alterations of neuronal morphology, such as neuronal projection, mediated by Rho GTPases, ultimately facilitating optimal learning and memory (99)(100). We also propose that the lincRNAs ENSRNOT00000076905 and ENSRNOT00000088470 may have been involved in this process by regulating the expression of genes implicated in the Rho GTPase signalling pathway.

Another pathway associated with the fear extinction related lincRNAs was the *immune system*. In recent years, there has been a growing awareness of the importance of the immune system in supporting optimal CNS functioning (101) and the detrimental effects that a dysregulated immune system can evoke on neuronal functioning (102)(103) and mental health (104)(105). Sufficient immune functioning not only supports optimal stress-coping responses, but is also essential for learning and memory (101)(102)(103). In this study, DCS may have facilitated fear extinction by regulating the expression of immune-related genes via lincRNAs such as ENSRNOT00000080023 and ENSRNOT00000076905.

As expected, the main molecular functions associated with fear extinction related lincRNAs were nucleotide, ribonucleotide and cytoskeletal protein binding, since the main features of lincRNAs are the regulation of gene and protein expression through its interactions with chromatin and RNAs or by recruiting and interacting with transcriptional repressors or enhancers (as reviewed by (106)). LncRNAs can even interact with DNA and one mechanism involved in direct RNA–DNA interactions are triple helices. Double-stranded DNA forms triple-helical structures by incorporating a third single-stranded nucleic acid in its major groove, forming Hoogsteen or reverse Hoogsteen hydrogen bonds with a purine-rich strand of DNA (107). Interestingly, other enriched molecular functions included purine nucleotide-binding and purine ribonucleotide binding. In the nucleus, these triple-helical structures (containing ribosomal DNA [rDNA] and lincRNA) are recognized by DNA methyltransferase DNMT3B, which methylates rDNA promoters and subsequently represses rDNA transcription (108). Moreover, certain lncRNAs directly interact with DNA in a sequence-specific manner and subsequently activates (109)(110) or repress transcription (111)(112) through the recruitment of coactivator or corepressor proteins. Some lncRNAs can also form triple helices in *cis* (auto-binding) (111)(110)(113), therefore enabling regulation of the exact locations they are transcribed from. Our findings therefore not only highlight the main functions of lincRNAs but also point to directions for future research, namely interrogation of lincRNA-DNA interactions and their subsequent effects on transcriptional and translational regulation.

LncRNAs can regulate the expression of neighbouring genes in-*cis* (38)(39). We therefore identified QTLs that overlapped with the genomic regions of four fear extinction related lincRNAs for which there was an RGD ID available (ENSRNOT00000081704, ENSRNOT00000084136, ENSRNOT00000088478 and ENSRNOT00000086613). The selected QTLs of interest were involved in traits such as serum corticosterone level, neuroinflammation as well as anxiety, stress and despair related responses. Dysregulation of the hypothalamic-pituitary-adrenal (HPA) axis results in an inability to initiate a normal stress response, which is a key feature of PTSD. The HPA axis is regulated by a negative feedback mechanism, where excess cortisol (or corticosterone in rodents) binds to glucocorticoid receptors in the hypothalamus and pituitary and subsequently suppresses the release of corticotropin-releasing hormone and adrenocorticotropin hormone. The HPA-axis also interacts with the immune system to maintain homeostasis (Wong et al., 2002) and there is an intricate relationship between the immune system, brain and behaviour, as discussed earlier. Research has also shown that immune functioning is affected in PTSD patients (114)(115)(116). We, therefore, hypothesise that the upregulation of these lincRNAs, following DCS administration, may result in the *cis*-regulation of genes that control cortisone levels and neuroinflammation, which elicited downstream effects on learning and memory to ultimately alleviate anxiety and stress-related responses and promote successful fear extinction.

## Conclusion

This study employed bioinformatics, *in silico* interaction prediction and gene set enrichment analyses to identify differentially expressed lincRNAs that may have targeted and regulated the expression of mRNAs that are enriched in biological processes, molecular functions and pathways that mediate fear extinction. Future studies could build on the *in silico* results by using cell-culture based assays to functionally verify predicted lincRNA-mRNA interactions. Protein quantification of predicted mRNA targets should also be performed; unfortunately, due to the limited quantity of hippocampal tissue, we could not determine whether changes in gene expression tranlsated to altered protein expression in this model.

This is the first study to identify lincRNAs and their RNA targets that play a role in transcriptional regulation during fear extinction. Our research identified differentially expressed lincRNAs and their predicted mRNA and pre-mRNA targets that could help us decipher the molecular basis of DCS-induced fear extinction. Four hippocampal lincRNAs, ENSRNOT00000076905, ENSRNOT00000088470, ENSRNOT00000080023 and ENSRNOT00000092675, interacted with nucleotides, ribonucleotides and proteins, thereby regulating the expression of genes involved in neuronal projection and neurogenesis, a dynamic process required during learning and memory, which was possibly mediated by the Rho GTPase pathway. Through the regulation of serum corticosterone levels, and subsequently, the HPA-axis, lincRNAs ENSRNOT00000081704, ENSRNOT00000084136, ENSRNOT00000088478 and ENSRNOT00000086613 may also have attenuated anxiety, stress and despair related responses through improved neuro-immune functioning. These eight lincRNAs were important role players in a well orchestrated sequence of events that resulted in effective fear extinction in this model investigating the core phenotypes of PTSD.

## Supporting information

all differentially expressed transcripts

differentially expressed mRNAs_FSMvsCS

differentially expressed mRNAs_FDWvsFSM

differentially expressed lincRNAs_FSMvsCS

differentially expressed lincRNAs_FDWvsFSM

fear_extinction_mRNAs_opposite

fear_extinction_lincRNAs_opposite

lincRNAs_interact_mRNAs

lincRNAs_interact_pre-mRNAs

Disease_enrichment

Molecular Functions

Biological_Pocesses

differentially expressed mRNA_FDW_FSM

differentially expressed mRNA_FSM_CS

differentially expressed lincRNA_FDW_FSM

differentially expressed lincRNA_FSM_CS

## Acknowledgements

This work is based on research supported by the South African Research Chairs Initiative of the Department of Science and Technology and National Research Foundation of South Africa, as well as the South African Medical Research Council (SAMRC) under a Self Initiated Research Grant. Thank you to Dr Fairbairn-Adonis for performing the animal work. A special thanks to Dr Oakeley and Novartis Pharma (Basel) for performing the RNA sequencing. MDR acknowledges support from the University Research Priority Program Evolution in Action at the University of Zurich. VBCS acknowledges support from the Brazilian institution *Conselho Nacional de Desenvolvimento Científico e Tecnológico* (CNPq).

