## Supplementary figures and images for "LincRNAs involved in DCS-induced fear extinction: Shedding light on the transcriptomic dark matter"

### differentially expressed lincRNA_FDW_FSM

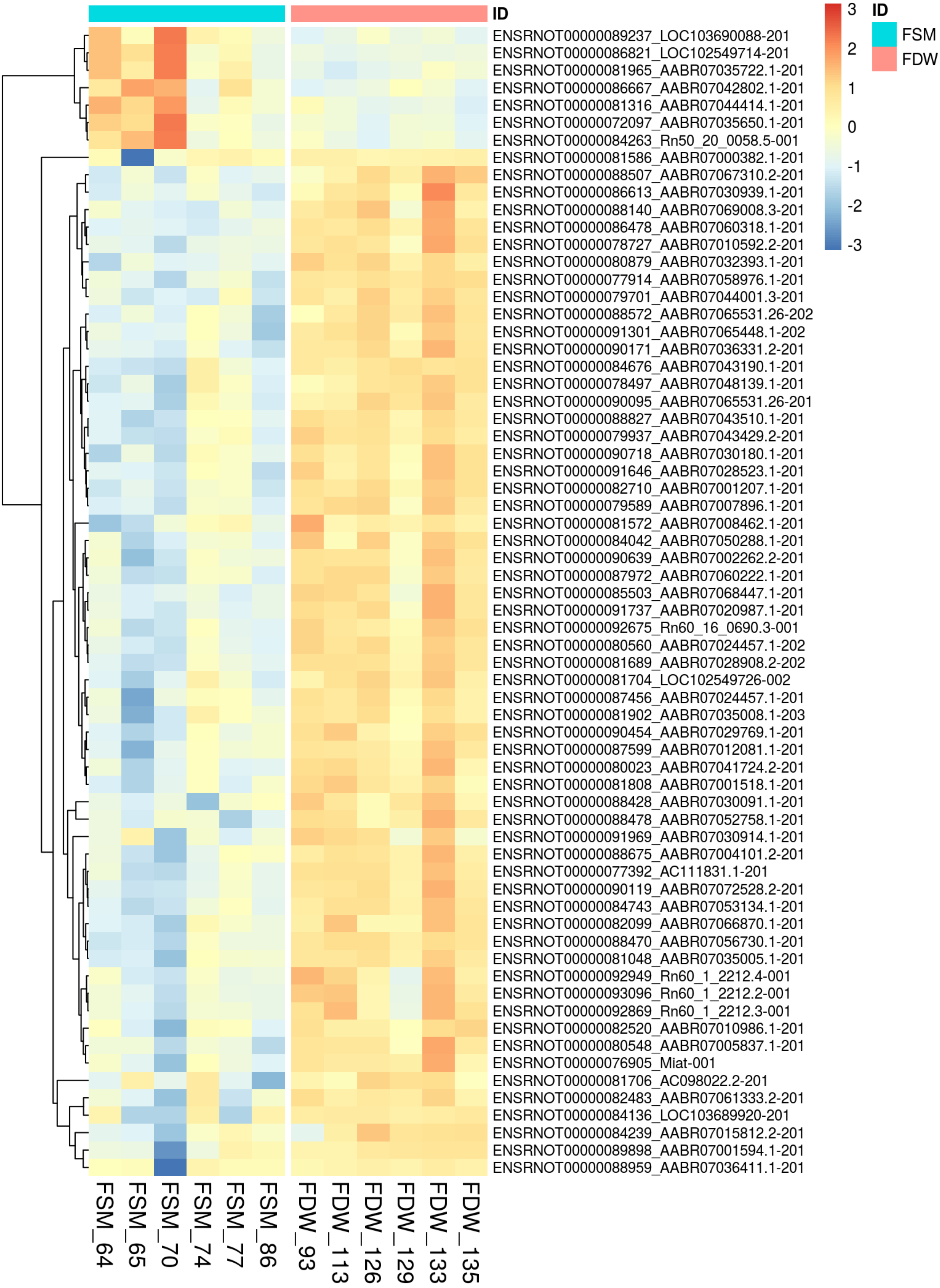

### differentially expressed lincRNA_FSM_CS

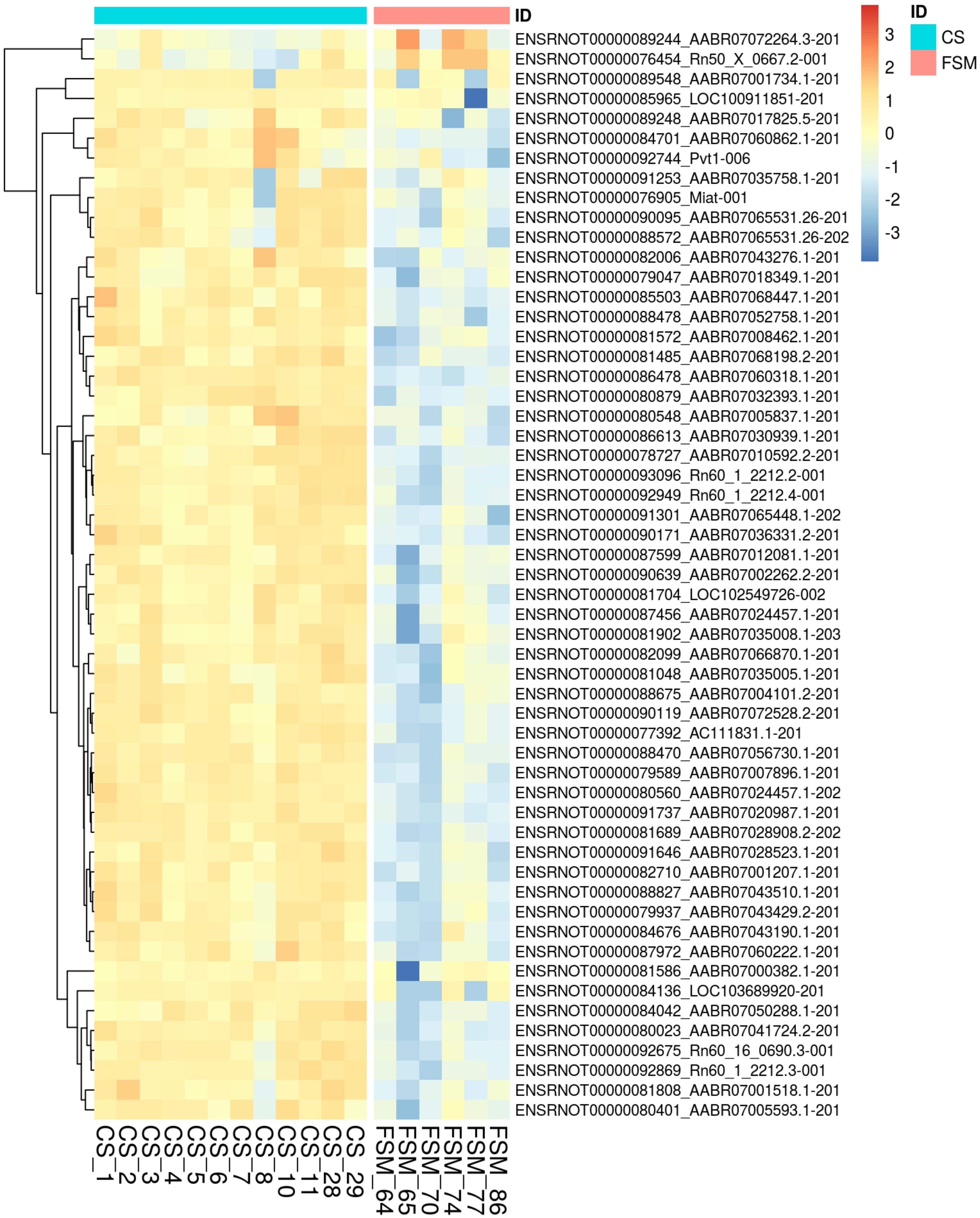

### differentially expressed mRNA_FDW_FSM

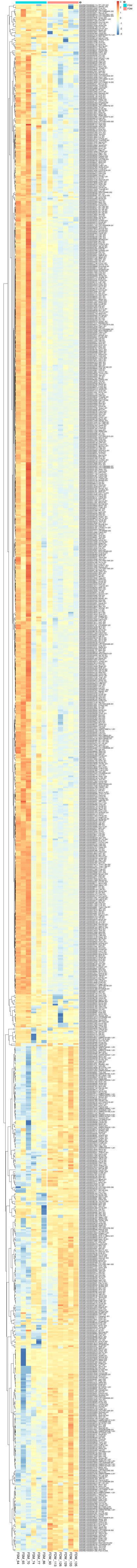

### differentially expressed mRNA_FSM_CS

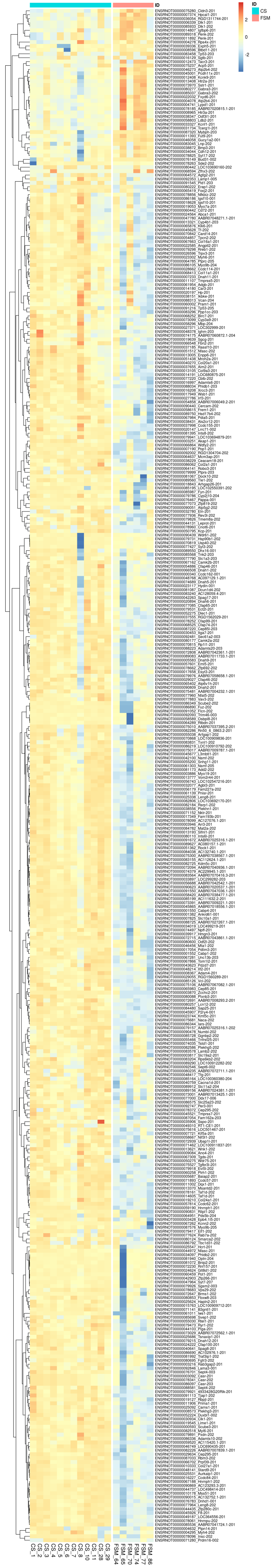
